# The AID2 system offers a potent tool for rapid, reversible, or sustained degradation of essential proteins in live mice

**DOI:** 10.1101/2024.06.04.597287

**Authors:** Valentina C Sladky, Margaret A Strong, Daniel Tapias-Gomez, Cayla E Jewett, Chelsea G Drown, Phillip M Scott, Andrew J Holland

## Abstract

Studying essential genes required for dynamic processes in live mice is challenging as genetic perturbations are irreversible and limited by slow protein depletion kinetics. The first-generation auxin-inducible-degron (AID) system is a powerful tool for analyzing inducible protein loss in cultured cells. However, auxin administration is toxic to mice, preventing its long-term use in animals. Here, we use an optimized second-generation AID system to achieve the conditional and reversible loss of the essential centrosomal protein CEP192 in live mice. We show that the auxin derivative 5-Ph-IAA is well tolerated over two weeks and drives near-complete CEP192-mAID degradation in less than one hour *in vivo*. Prolonged CEP192 loss led to cell division failure and cell death in proliferative tissues. Thus, the second-generation AID system is well suited for rapid and/or sustained protein depletion in live mice, offering a valuable new tool for interrogating protein function *in vivo*.

## Introduction

Loss-of-function mutant mice are critical parts of the genetic toolbox for basic research. However, DNA and RNA-based knock-out and knock-down approaches have limitations, including irreversibility and slow depletion dynamics. These shortcomings can be overcome by induced degradation with chemical methods such as PROTACs (Proteolysis Targeting Chimeras) (*1*), or chemical genetic approaches like the dTag system or AID system (*2–4*). PROTACs are bivalent binders that bridge the protein of interest and an E3 ligase to promote proximity-induced ubiquitylation (*5*, *6*). PROTACs require the engineering of chemical ligands that specifically bind the target protein, limiting their applications to ligandable targets. The dTag system requires tagging the gene of interest with an FKBP12^F36V^ tag so that a heterobifunctional degrader can recruit the FKBP12^F36V^-tagged target to CRBN for ubiquitination and degradation (*7*). dTag has proven valuable for short-term applications, achieving near-complete protein depletion in xenograft tumors, adult tissues, and embryos (*8–10*). However, long-term applications in mice are limited by the toxicity of the vehicle and the small molecule required to induce protein degradation (*11*). In addition, PROTACs and the dTAG system are based on heterobifunctional molecules that can display a Hook effect due to the saturated binding of the ligase and degradation target at high concentrations (*12*).

The auxin-inducible-degron (AID) system utilizes the plant hormone auxin, which acts as a molecular glue to stabilize the binding of an auxin-inducible-degron to the plant-derived E3 ligase adaptor Tir1 (transport inhibitor response 1). In the presence of the auxin hormone indole-3-acetic acid (IAA), the TIR1-SCF E3 ligase complex targets AID-tagged proteins for degradation (*13*). The AID system has been applied successfully in mammalian cells to drive inducible protein degradation, reducing the half-life of targeted proteins to < 30 minutes (*13*, *14*). However, the original AID system suffers from leakiness and requires high IAA concentrations (100-500 µM). These limitations were overcome with the development of a second-generation AID system (AID2), in which a point mutation in *Tir1* (F74A or F74G) promotes an interaction with the bulky IAA derivatives 5-Adamantyl-indole-3-acetic acid (5- Ad-IAA) or 5-phenyl-indole-3-acetic acid (5-Ph-IAA) at up to 1000 fold lower concentrations (*15*). The AID2 system has been shown to work for *in vivo* applications in *D. melanogaster* (*16*) and *C. elegans* (*17*).

Recent efforts have focused on applying the AID system for *in vivo* use in mice (*18–20*). The first-generation AID system has been used to target CDC7 (*20*) and the condensin subunits NCAPH and NCAPH2 (Macdonald *et al.*, 2022) for degradation in mice expressing *Oryza sativa Tir1 (OsTir1)* from the *Rosa26* locus. Both studies report protein depletion within 2-6 hours in the cell types tested (*19*, *20*). However, *in vivo* IAA treatment caused significant toxicity in mice; therefore, most experiments were performed *ex vivo* (*19*, *20*). A recent proof-of-principle study tested the second-generation AID system in mice expressing a randomly integrated *OsTir1*-F74G transgene. Injections of the auxin derivative 5-Ph-IAA achieved robust protein degradation within 6 hours in various tissues. However, these results were limited to analyzing the degradation of a randomly integrated GFP reporter carrying an AID tag, and neither long-term 5-Ph-IAA treatments nor pathological assessments were performed (*18*).

The AID system has been widely used in cultured cell lines to study essential genes in cellular processes such as mitosis and centrosome biology (*14*, *21–24*); however, it has not yet been applied to analyzing essential genes in mice. CEP192 is a centrosomal protein characterized as common essential gene by the Cancer Dependency Map (Depmap). The centrosome is a membrane-less organelle consisting of a pair of centrioles surrounded by pericentriolar material (PCM) and functions as a microtubule organizing center in some interphase cells. Centrioles are duplicated once during S-phase to ensure their numbers are maintained from one generation to the next. During mitosis, the centrosome undergoes maturation and expands the PCM to increase microtubule nucleation and form the bipolar mitotic spindle (*25*). In quiescent cells, modified centrioles function as basal bodies that template cilia critical for signaling, fluid transport, and locomotion (*26*). CEP192 is essential for centriole duplication (*27–29*) and centrosome maturation to establish the bipolar mitotic spindle apparatus (*30–32*). Whether CEP192 is also directly involved in ciliogenesis is not known. Almost all the analysis of mammalian centrosomes and CEP192 has been performed in cultured cells. However, recent work has shown that centrosome composition and function can vary across mammalian tissues and cell types (*33–36*). Moreover, analyzing CEP192 function in mouse models *in vivo* is challenged by the fast dynamics of centriole duplication and cell division. Here, we utilize the second-generation AID system in live mice to investigate the *in vivo* functions of the essential centrosome protein CEP192.

## Results

### Creation of CEP192-mAID-mNeonGreen and conditional OsTir1-F74G mouse lines

To evaluate the utility of the first-generation AID system in mice, we first examined the effects of administering IAA at doses used previously to achieve protein degradation in mice. Intraperitoneal injection of a single dose of IAA at 400 mg/kg or 800 mg/kg in PBS induced spasms and paralysis within 1 hour or 15 minutes, respectively (Fig. 1A, S1A-B). While some mice dosed with 400 mg/kg recovered, all animals receiving 800 mg/kg reached the humane endpoint with near-complete paralysis after one hour (Fig. 1B, S1A). We conclude that IAA is toxic in mice at the doses required to induce protein degradation (Fig. S1C) and thus, the first- generation AID system is unsuitable for the *in vivo* manipulation of protein levels.

**Figure 1:**
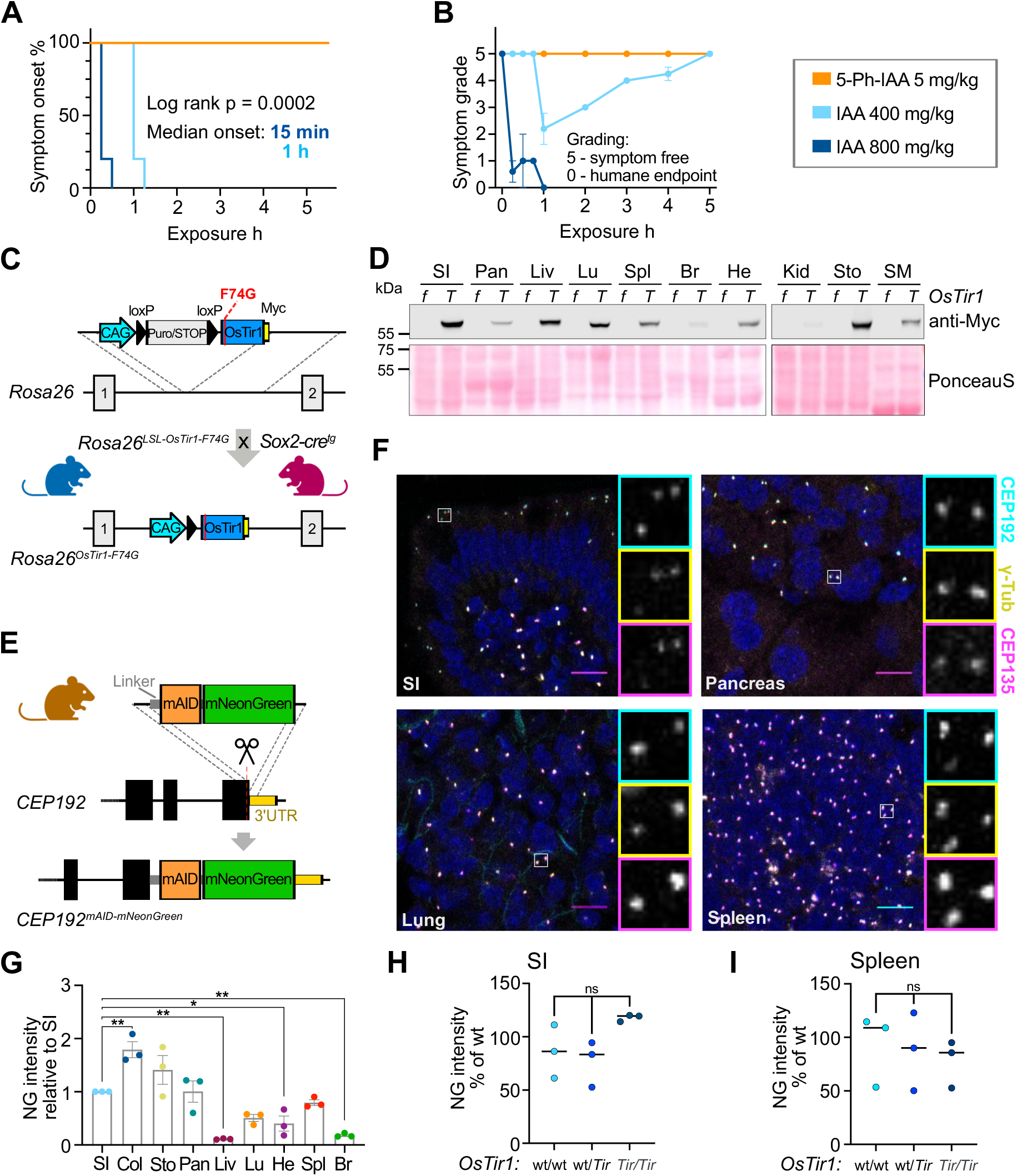
**Characterization of OsTir1-F74G and Cep192^mAID-mNG^ mice.** (A) Kaplan-Meier-plot showing symptom onset after i.p. injection of OsTir1^Tir/Tir^ mice with 5mg/kg 5-Ph-IAA (N = 3), 400mg/kg IAA (N = 5), or 800mg/kg IAA (N = 5). (B) Symptoms including spasms and paralysis were graded by severity, 0 = symptom free, 5 = near complete paralysis, humane endpoint. 5mg/kg 5-Ph-IAA (N = 3), 400mg/kg IAA (N = 5), or 800mg/kg IAA (N = 5). (C) Schematic showing the targeting strategy to integrate the lox-stop-lox (LSL)-OsTir1-Myc construct at the Rosa26 locus. (LSL)-OsTir1-Myc animals were crossed to Sox2-Cre expressing mice to achieve whole-body expression of OSTIR1-F74G-Myc. (D) Immunoblot probed with an antibody detecting Myc-tagged OSTIR1 in the noted organs isolated from an OsTir1^f/f^ (f) and an OsTir1^Tir/Tir^ (T) mouse. SI – small intestine, Pan – Pancreas, Liv – Liver, Lu – Lung, Spl – Spleen, Br – Brain, He – Heart, Kid – Kidney, Sto – Stomach, SM – Skeletal Muscle. Ponceau-S-staining is shown as a loading control. A biological replicate with a second pair of genotypes is shown in Supplemental Fig. 1G. (E) Schematic illustration showing the strategy to target exon 48 of Cep192 to endogenously tag the C-terminus of Cep192 with mAID and moxNeonGreen (Cep192^AID^). (F) Representative confocal immunofluorescence images of the indicated organs showing CEP192^AID^ signal (NeonGreen; cyan), and immunostained γ-tubulin (yellow) and CEP135 (magenta); Scale bar 10 µm. (G) Quantification of the CEP192^AID^ NeonGreen (NG) signal intensity in the noted tissues was measured by fluorescence microscopy and is shown relative to the small intestine (SI). Col – Colon, Sto – Stomach, Pan – Pancreas, Liv – Liver, Lu – Lung, He – Heart, Spl – Spleen, Br – Brain. N = 3 mice. (H-I) Graphs showing the NeonGreen (NG) signal quantified in the SI (H) and the spleen (I) of untreated Cep192^AID^ mice that were either wt/wt, wt/Tir, or Tir/Tir for OsTir1. N = 3 per genotype. Data is shown as mean ± SEM. Statistical significance was determined using one-way ANOVA with Sidak’s multiple comparisons test (G-I). ns p≥0.05, * p<0.05, ** p<0.01.

We next analysed the effect of the IAA derivative 5-Ph-IAA that is used in the AID2 system at similar doses to those used previously in mice (*18*). Mice injected with 5 mg/kg 5-Ph-IAA in PBS were symptom-free over the observation period of 5 hours (Fig. 1A-B, S1A). This motivated us to develop mouse models to test the second-generation AID system *in vivo*. We created a conditional allele of *OsTir1-F74G* by targeting a loxP-Stop-loxP cassette followed by *OsTir1-F74G-Myc* to the *Rosa26* locus. The resulting *Rosa26-OsTir1-F74G^flox^* mice were crossed to *Sox2-Cre* animals that express Cre in the germline to generate *Rosa26-OsTir1-F74G^Tir^* animals, hereafter referred to as *OsTir1^Tir^* (Fig. 1C). *OsTir1^Tir^* animals had a similar body weight to *OsTir1^flox^* mice (Fig. S1D). Homozygous *OsTir1^Tir/Tir^* mice were fertile, and heterozygous breedings produced offspring at Mendelian ratios (Fig. S1E-F). Importantly, the OSTIR1 protein was expressed across various tissues, with the lowest expression detected in the brain and kidney (Fig. 1D, S1G).

To test the effectiveness of *OsTir1*-F74G mediated protein degradation, we endogenously tagged the C-terminal exon of *Cep192* with a flexible linker, a miniaturized AID tag (mAID, (*37*)) and moxNeonGreen to track protein abundance (Fig. 1E). Heterozygous and homozygous *Cep192^mAID-moxNeonGreen^* mice (hereafter *Cep192^AID^*) are fertile and produce offspring at the expected Mendelian ratios (Fig. S1E, H). Body weights of the various combinations of *Cep192^AID^; OsTir1^Tir^* genotypes were similar to *OsTir1^flox^* mice (Fig. S1D), indicating that tagging did not interfere with the essential roles of CEP192. Immunofluorescence analysis showed that endogenously tagged CEP192 was expressed across various tissues and co-localized with the centrosomal proteins CEP135 and γ-tubulin (Fig. 1F-G, S1I-J). While CEP192^AID^ abundance at the centrosome was similar across cell types in most tissues, liver hepatocytes displayed minimal CEP192^AID^ expression and CEP192^AID^ was only detected in non-parenchymal liver cells (Fig. S1K). Leaky degradation was not observed in the absence of 5-Ph-IAA in the small intestine (SI) and spleen of *Cep192^AID/AID^* animals heterozygous or homozygous for *OsTir1^Tir^* (Fig. 1H-I). Together, we conclude that 5-Ph-IAA can overcome the toxicity-related limitations of IAA making the AID2 system suitable for use *in vivo*. Moreover, we find that the homozygous *OsTir1*-*F74G* and *Cep192^AID^* alleles are well tolerated and widely expressed in mice.

### CEP192^AID^ is rapidly degraded in primary cells

To analyse the dynamics of CEP192^AID^ degradation we generated *Cep192^AID/AID^* MEF lines from embryos wt/wt, *Tir*/wt, or *Tir*/*Tir* for *OsTir1* and measured the abundance of CEP192^AID^ at the centrosome using immunofluorescence. Treatment with 1 µM 5-Ph-IAA led to near-complete degradation of centrosomal CEP192^AID^ within 1 hour in both heterozygous and homozygous *OsTir1* lines (Fig. 2A-B). The degradation maximum (D_max_) was 98% for wt/*Tir* and 97% for *Tir*/*Tir* genotypes (Fig. 2A-B). We further analysed CEP192^AID^ degradation dynamics using live imaging and confirmed efficient degradation of CEP192^AID^ within 1 hour in both *OsTir1* homozygous and heterozygous cells (Fig. S2A). Co-staining with an antibody raised against murine CEP192 confirmed complete protein degradation at 5 hours after 5-Ph-IAA addition (Fig. S2B). Consistent with previous reports, the abundance of the centrosome protein γ- tubulin in interphase was not significantly impacted by CEP192 degradation in cells (Fig. S2B) (*30*). However, 5-Ph-IAA treated *Cep192^AID/AID^* MEF cells expressing OSTIR1 almost exclusively formed monopolar spindles during mitosis (Fig. 2B-C) and live imaging showed that over 80% of MEFs lacking CEP192 delayed in mitosis and failed to divide before re-adhering to the plate (Fig. 2D-E, Fig S2C). Thus, CEP192^AID^ can be effectively degraded using the AID2 system in MEFs revealing a strong requirement of CEP192 for successful cell division.

**Figure 2:**
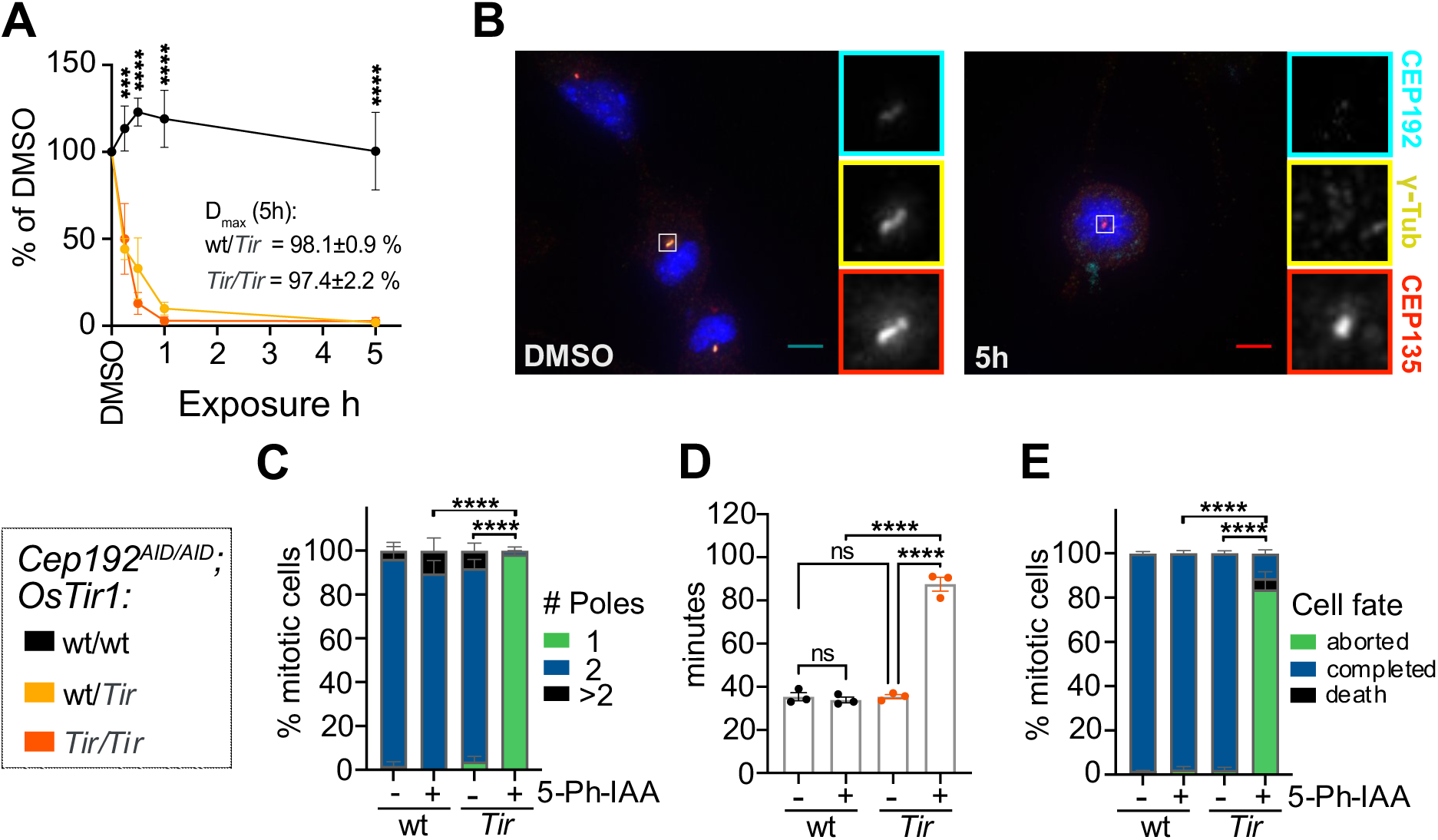
**The AID2 system efficiently degrades CEP192^AID^ in primary cultures.** (A) Quantification of NeonGreen (NG) intensity in immunofluorescence microscopy images relative to DMSO in MEF lines of the indicated genotypes treated with 5-Ph-IAA for various times. N = 3-5 MEF lines per genotype. (B-C) *Cep192^AID/AID^* MEFs that were either wt/wt or *Tir/Tir* for *OsTir1* were treated with 5-Ph-IAA (+) or DMSO (-) for 5h. (B) Representative images of OSTIR1 expressing MEFs immunostained for γ-tubulin (yellow) and CEP135 (red). CEP192^AID^ is shown in cyan. Insets show centrosomes in mitotic cells. Scale bar 5 µm. (C) Graph showing the number of mitotic spindle poles quantified in immunofluorescence images as in (B); n = 10-30 mitotic cells in N = 4 MEF lines per genotype. (D-E) *Cep192^AID/AID^* MEFs with or without OSTIR1 expression were treated with 5-Ph-IAA (+) or DMSO (-) and live imaged to determine the duration and fate of mitosis. (D) Quantification of the time spent in mitosis from rounding up until completion of cell division or cell death. (E) Graph showing the cell fate after mitosis: cells completed mitosis by successful division (blue), died (black), or re-adhered without division (green). N = 4 MEF lines per genotype. All data is displayed as mean ± SEM. Statistical significance was determined using one-way (D) or two-way (A, C, E) ANOVA with Sidak’s multiple comparisons test. ns p≥0.05, *** p<0.001, **** p<0.0001.

### CEP192 is not required for the formation or maintenance of primary or motile cilia

The requirement of CEP192 for centriole duplication has prevented a clear analysis of its role in ciliogenesis. We therefore exploited the rapid depletion of CEP192^AID^ in primary MEFs to investigate CEP192’s function in ciliogenesis. *Cep192^AID/AID^* MEF lines with or without *OsTir1* were serum starved for 24 hours in the presence of 5-Ph-IAA. The number of cells with a primary cilium was unaffected by CEP192^AID^ degradation (Fig. 3A-B, Fig. S3A). Similarly, degradation of CEP192^AID^ for 24 or 48 hours after the primary cilium was established did not alter the number of ciliated cells (Fig. S3B-C). We conclude that CEP192 is dispensable for the formation and maintenance of the primary cilium.

**Figure 3:**
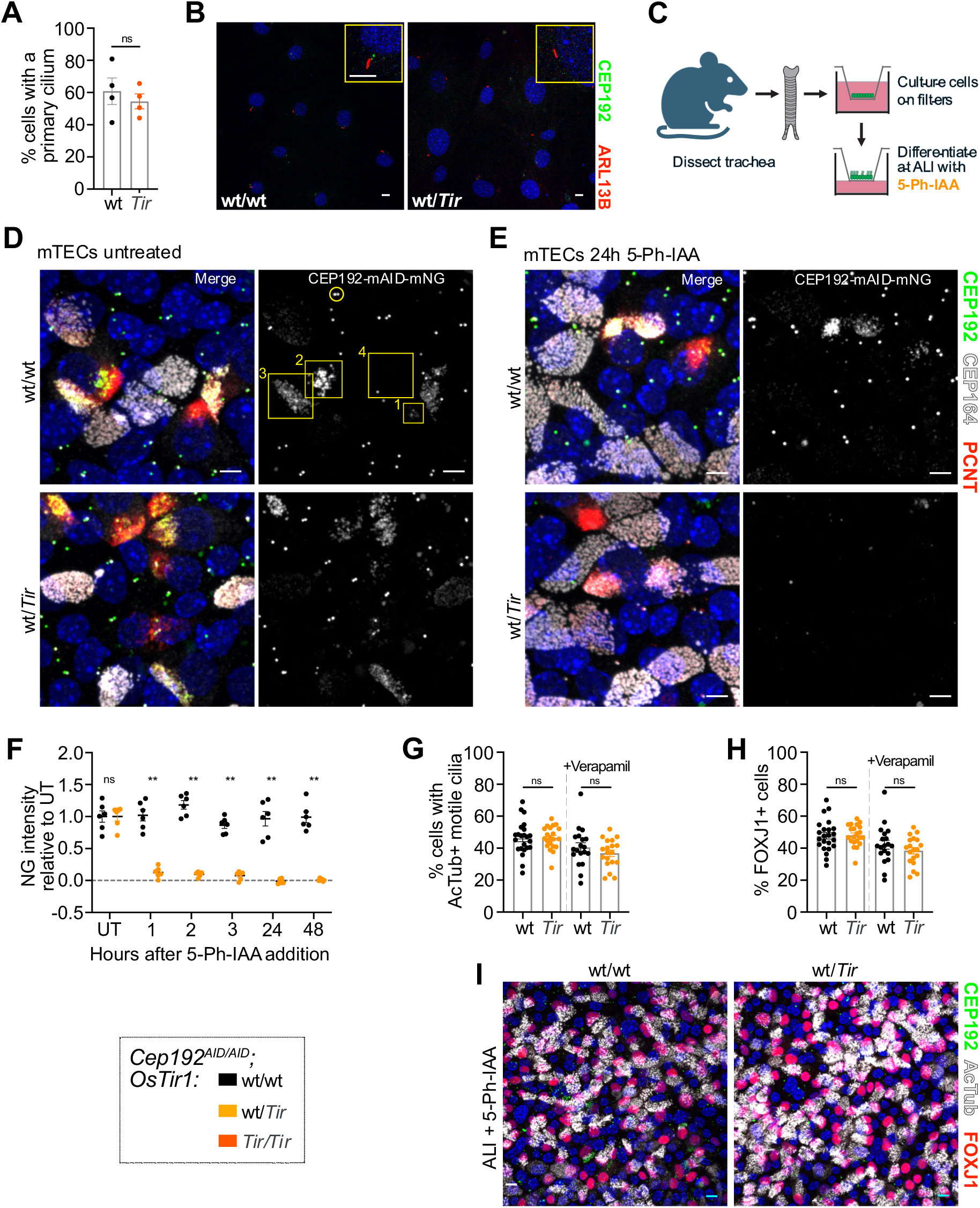
**CEP192 is not required for primary or motile cilia** (A-B) MEF lines of the indicated genotypes were serum starved in the presence of 5-Ph-IAA for 24h. (A) Quantification of the number of cells with a primary cilium; N = 2 MEF lines per genotype from 2 separate passages. (B) Representative confocal image of MEFs expressing CEP192^AID^ NeonGreen (green), and immunostained for the primary cilia marker Arl13B (red). Yellow boxes show insets of a primary cilium. Scale bar 5 µm. (C) Schematic illustration showing isolation and differentiation of mTECs. 5-Ph-IAA was added to the basal media of air-liquid-interphase cultures (ALI). (D) Representative confocal images of mTECs with (wt/*Tir*) or without (wt/wt) OSTIR1 expression at ALI day 5 expressing CEP192^AID^ (green) and immunostained for PCNT (red) and CEP164 (gray). Yellow circle marks the CEP192^AID^ signal at the centrosome in non-differentiating cells. Yellow boxes mark the CEP192^AID^ signal in differentiating cells with numbers denoting early (1), mid (2), and late (3), and complete (4) differentiation. Scale bars 5 µm. (E) Representative confocal images of mTECs at ALI day 5 were treated with 5-Ph-IAA for 24h. CEP192^AID^ signal is shown in green and tissues were immunostained for PCNT (red) and CEP164 (gray). Scale bars 5 µm. (F) Time course showing CEP192^AID^ intensity at newly amplified centrioles following 5-Ph-IAA treatment in differentiating cells without (wt/wt) or with OSTIR1 expression (wt/*Tir*). The signal intensity was normalized to the untreated (UT) condition. N = 2 mice per genotype indicated by different symbols, n=6 fields of view. (G-I) mTECs expressing CEP192^AID^ either wt or heterozygous for *OsTir1* were treated with 5-Ph-IAA for ALI days 0-7. Cells were fixed on ALI day 7 and immunostained for the differentiation marker FOXJ1 and for acetylated tubulin to label motile cilia (AcTub). (G) Graph showing the percentage of cells with motile cilia in mTEC cultures in the absence or presence of Verapamil. (H) Quantification of the percentage of cells in mTEC cultures with FOXJ1+ nuclei with or without Verapamil. (G-H) N = 3 mice per genotype indicated by different symbols, n=6 fields of view per genotype and condition. (I) Representative confocal image of mTEC cultures on ALI day 7 expressing CEP192^AID^ (green) immunostained for FOXJ1 (red) and AcTub (gray). Scale bars 5 µm. All data is displayed as mean ± SEM. Statistical significance was determined using two-way ANOVA with Sidak’s multiple comparisons test (F), or a two-tailed, unpaired Student’s t-test (A, G-H). ns p≥0.05, ** p<0.01.

To further evaluate the role of CEP192 in ciliogenesis we used *ex vivo* cultures of mouse tracheal epithelial cells (mTECs), which differentiate into multiciliated cells (MCCs) that amplify centrioles to build hundreds of motile cilia at their apical cell surface. To generate mTECs, tracheas were dissected from *Cep192^AID/AID^* mice with or without *OsTir1* and cultured on transwell filters to form a polarized epithelial monolayer. Cells were then differentiated at an air-liquid-interface (ALI) and 5-Ph-IAA was added to media in the basal chamber at various time points (Fig 3C). The resulting culture contains several cell types, including MCCs. CEP192^AID^ localized to centrosomes of non-MCCs (Fig 3D, yellow circle) and newly born, amplified centrioles in MCCs (Fig 3D, yellow boxes). In early stages of MCC differentiation, the CEP192^AID^ signal clustered around the centrioles and decreased as cells completed differentiation with motile cilia (Fig 3D, yellow box labeled 4). After 1 hour of 5-Ph-IAA treatment, CEP192^AID^ was nearly undetectable in MCCs and non-MCCs in the presence of OSTIR1, and stayed low over the entire observation period of 48 hours (Fig. 3D-F, Fig. S3D).

Given that CEP192 localizes to centrioles during MCC differentiation, we next tested whether CEP192 was required for centriole amplification and motile cilia formation. *Cep192^AID/AID^* mTECs with or without *OsTir1* were cultured at ALI for 7 days in the presence of 5-Ph-IAA. We observed no difference in the number of MCCs (FOXJ1+ cells) or cells with motile cilia (Fig 3G-I). We also performed these experiments in the presence of verapamil, an efflux pump inhibitor, as some drugs are ineffective in mTECs due to the efflux capability of these epithelial cells (*38*). 5-Ph-IAA effectiveness was not impacted by verapamil (Fig 3G-H), suggesting that this compound is not sensitive to efflux pumps. To assess whether CEP192 is required for cilia maintenance, we allowed the cultures to differentiate for 7 days at ALI and then treated with 5-Ph-IAA for 2 days (ALI days 7-9). The number of FOXJ1+ cells and cells bearing motile cilia was similar, showing CEP192 is dispensable for cilia maintenance (Fig. S3E-G). Taken together, we conclude CEP192 function is not required for differentiation, centriole amplification, or motile cilia formation and maintenance in MCCs.

### AID2 induced rapid and reversible CEP192^AID^ degradation *in vivo*

We next examined the *in vivo* degradation dynamics of CEP192^AID^ in different tissues. *Cep192^AID/AID^*; *OsTir1^Tir/Tir^* mice were injected intraperitoneally with 5 mg/kg 5-Ph-IAA in PBS and CEP192^AID^ signal intensity (NeonGreen) was analyzed by immunofluorescence at various time points. 30 minutes after 5-Ph-IAA administration, CEP192^AID^ was reduced by ∼90% in the small intestine (SI) and > 80% in the spleen and the stomach (Fig. 4A-E, I, Fig. S4A). At the same time point, ∼70% of CEP192^AID^ was degraded in the pancreas and lungs, and 60% was degraded in non-parenchymal liver cells (Fig. 4F-I, Fig. S4A). D_max_ reached 94% in the spleen and 98% in the SI 5 hours after 5-Ph-IAA injection (Fig. 4A-B). The CEP192^AID^ signal remained low for 24 hours after 5-Ph-IAA administration and showed almost complete recovery by 72 hours post-injection in both tissues (Fig. 4A, J). We also tested the effect of a lower 1 mg/kg dose of 5-Ph-IAA (Fig. S4B-C). Treatment with 1 mg/kg degraded ∼80% of CEP192^AID^ in the SI but only reduced CEP192^AID^ by 53% in the spleen (Fig. S4B-C). The lower dose of 5-Ph-IAA also resulted in faster recovery of CEP192^AID^ in both tissues. Together, these data show the AID2 system allows reversible, near-complete degradation of CEP192^AID^ in less than 3 hours in different tissues, and degradation and recovery dynamics can be tuned by titrating the dose of 5-Ph-IAA.

**Figure 4:**
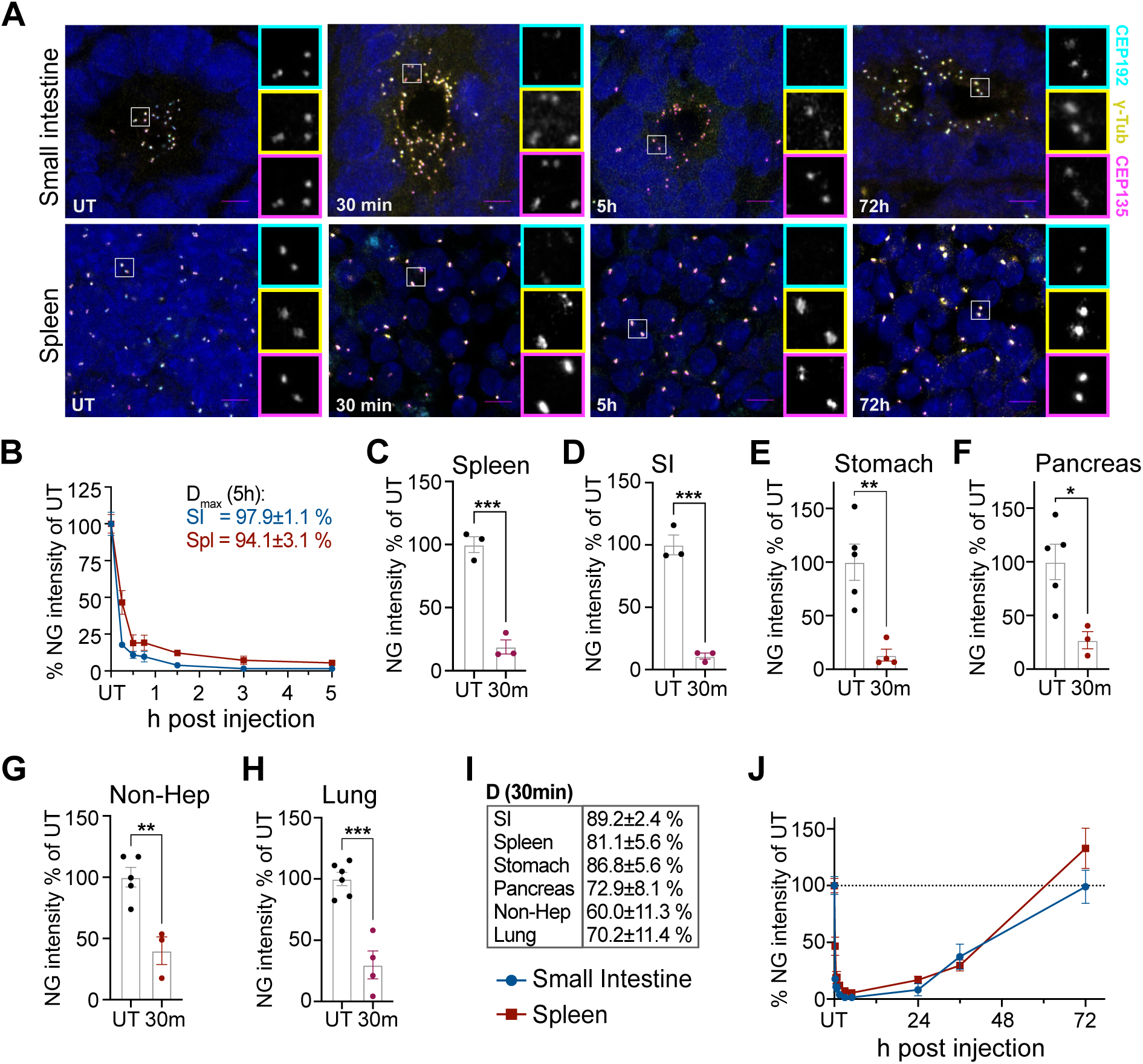
**The AID2 system induces rapid and reversible CEP192^AID^ degradation in mice.** (A) Representative confocal images of the small intestine and the spleen of *CEP192^AID^* mice expressing OSTIR1 that were left untreated (UT), or analysed at various times after i.p. injection of 5-Ph-IAA. CEP192^AID^ (cyan), γ-tubulin (yellow), CEP135 (magenta); scale bars 5 µm. (B) Graph showing the CEP192^AID^ intensity (NeonGreen, NG) in the small intestine (SI, blue) and the spleen (red) measured by fluorescence microscopy relative to the untreated (UT) control at 15 min, 30 min, 45 min, 1.5 h, 3 h, and 5 h after i.p. injection. Maximum degradation (Dmax) was reached by 5h. (C-H) Quantifications of the CEP192^AID^ (NeonGreen, NG) signal intensity in the indicated organs are shown relative to the untreated controls (UT) 30 min after treatment. (G) Since CEP192^AID^ was not expressed in hepatocytes, the signal was quantified in non-parenchymal cells in the liver (Non-Hep). (I) Table shows the degree of degradation at 30 min for the organs in (C-H). (J) The CEP192^AID^ signal (NG) was quantified at the indicated time points up to 72h after 5-Ph-IAA injection. Data points for UT and time points up to 5h are the same as in (B). The dashed line indicates 100%; N = 3-6 mice per time point. Data is shown as mean ± SEM. Statistical significance was determined using a two-tailed, unpaired Student’s t-test, * p<0.05, ** p<0.01, *** p<0.001, **** p<0.0001.

### The AID2 system is suitable for long-term repeat dose experiments in mice

Since the AID2 system proved highly effective at inducing acute CEP192^AID^ degradation, we next tested the impact of chronic 5-Ph-IAA administration in mice. We injected *Cep192^wt/wt^; OsTir1^Tir/Tir^* mice (n=6, 3f/3m) every 24 hours for 14 days with 5 mg/kg 5-Ph-IAA or vehicle control (PBS; Fig. 5A). Daily monitoring of animal behavior and body weight showed no abnormalities (Fig. 5B). After the last injection, a full necropsy was performed and over 40 tissues were evaluated by a board-certified veterinary pathologist. 5-Ph-IAA treatment did not affect any of the tissues examined. Complete blood cell counts and serum analysis were also indistinguishable between the PBS or 5-Ph-IAA treated groups (Fig. 5C-D, Fig. S5A-B). This shows that repeated injections with an effective concentration of 5-Ph-IAA are well tolerated over at least 14 days.

**Figure 5:**
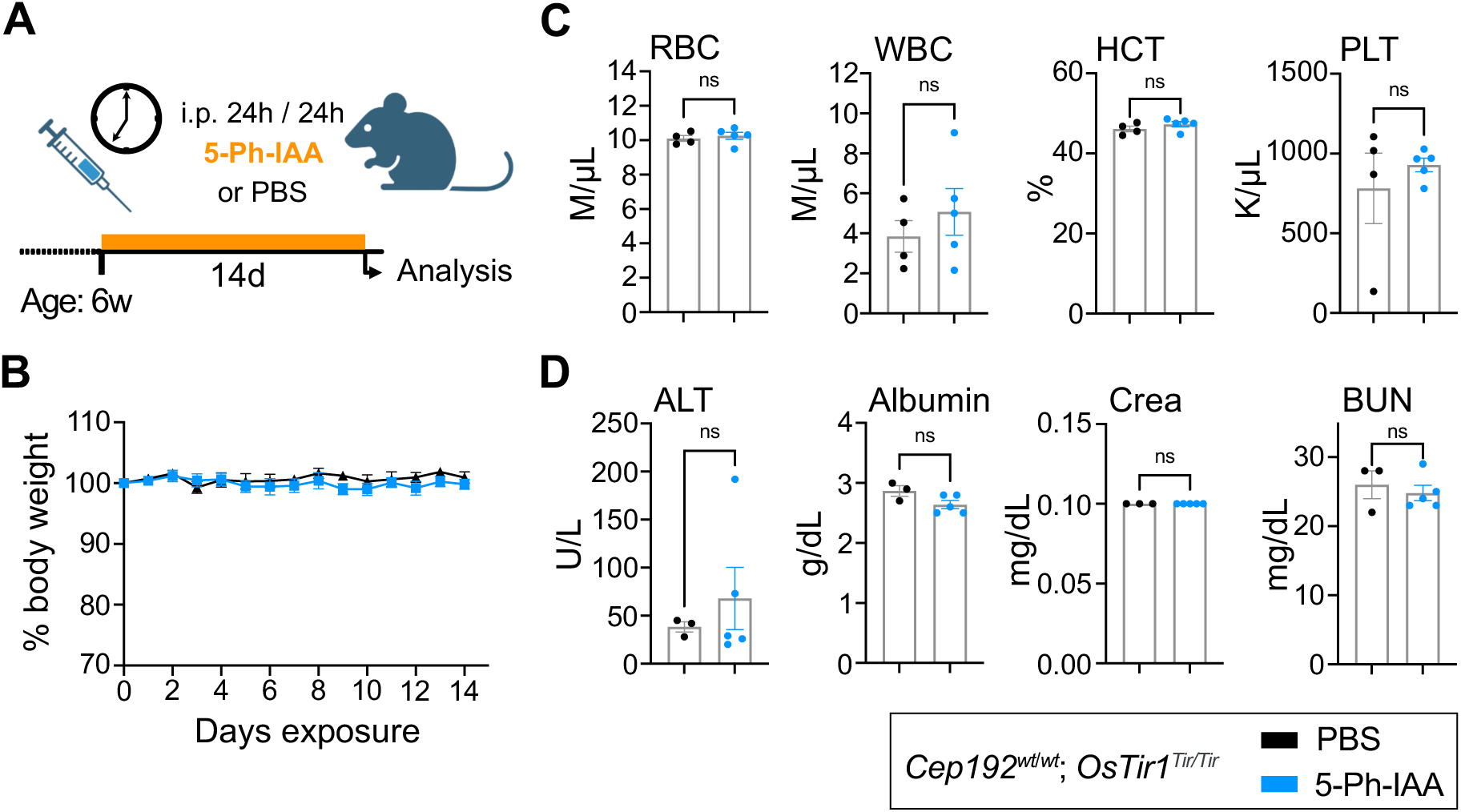
**Repeated dosing with 5-Ph-IAA is well tolerated over time.** (A) Schematic illustration of the treatment plan. 6-week-old *OsTir1^Tir/Tir^* mice were injected i.p. with 5-Ph-IAA or PBS every 24h for 14 days. (B) Graph showing body weight changes of mice treated as described in (A) and monitored over the treatment period. N = 6 mice per treatment. (C-D) Blood cell counts and serum parameters indicating toxicity were measured after 14 days of treatment. Full panels are shown in Supplemental Fig. 5. N = 6 mice per treatment. (C) Blood cell counts: RBC – red blood cells, WBC – white blood cells, HCT – Haematocrit, PLT – platelet count. (D) Serum parameters: ALT – alanine aminotransferase, Crea – Creatinine, BUN – blood urea nitrogen. Data is displayed as mean ± SEM. Statistical significance was assessed by a two-tailed, unpaired Student’s t-test; ns p≥0.05.

### Long-term CEP192 depletion causes weight loss and cell death in the spleen and intestine

Since 5-Ph-IAA was well-tolerated in *OsTir1^Tir/Tir^* mice, we investigated the long-term effects of depletion of CEP192 in mice. Since we observed minimal recovery of CEP192^AID^ after 24 hours (Fig. 4J), we injected *Cep192^AID/AID^*; *OsTir1^wt/Tir^* mice every 12 h for 3 or 8 days with 5-Ph-IAA or PBS (Fig. 6A). After 3 days, CEP192^AID^ protein level at the centrosome was reduced by > 95% in the spleen, SI, and other tissues analysed (Fig. 6B-D, Fig. S6A-C). Repeated 5-Ph-IAA dosing also reduced centrosomal γ-tubulin by half in the spleen and the SI at day 3 and this was maintained to the end of the experiment (Fig. 6B, E-F). As expected, repeated dosing with 5-Ph-IAA was well-tolerated in control mice, with minimal weight loss. By contrast, *Cep192^AID/AID^*; *OsTir1^wt/^*^Tir^ mice lost up to 20% body weight (humane endpoint) after 8 days of 5-Ph-IAA treatment (Fig. 6G). These animals displayed gastrointestinal symptoms and massive cell death in crypts of the small intestine and the colon, as indicated by active caspase-3 staining (Fig. 6H-i; Fig. S6D-G). While the level of cell death was not significantly elevated in the spleen following CEP192^AID^ loss (Fig. S6H-I), the spleens of *Cep192^AID/AID^*; *OsTir1^wt/Tir^* mice were markedly smaller after eight days of 5-Ph-IAA treatment (Fig. 6J). Importantly, CEP192 degradation resulted in a threefold increased mitotic index in the intestinal crypts and mitotic figures had a monopolar morphology, similar to our findings in MEFs (Fig. 2B-C, 6K-L). Together, these data show that repeated dosing with 5-Ph-IAA results in the degradation of >95% of centrosomal CEP192^AID^ *in vivo*, leading to mitotic delays and cell death in proliferating intestinal cells

**Figure 6:**
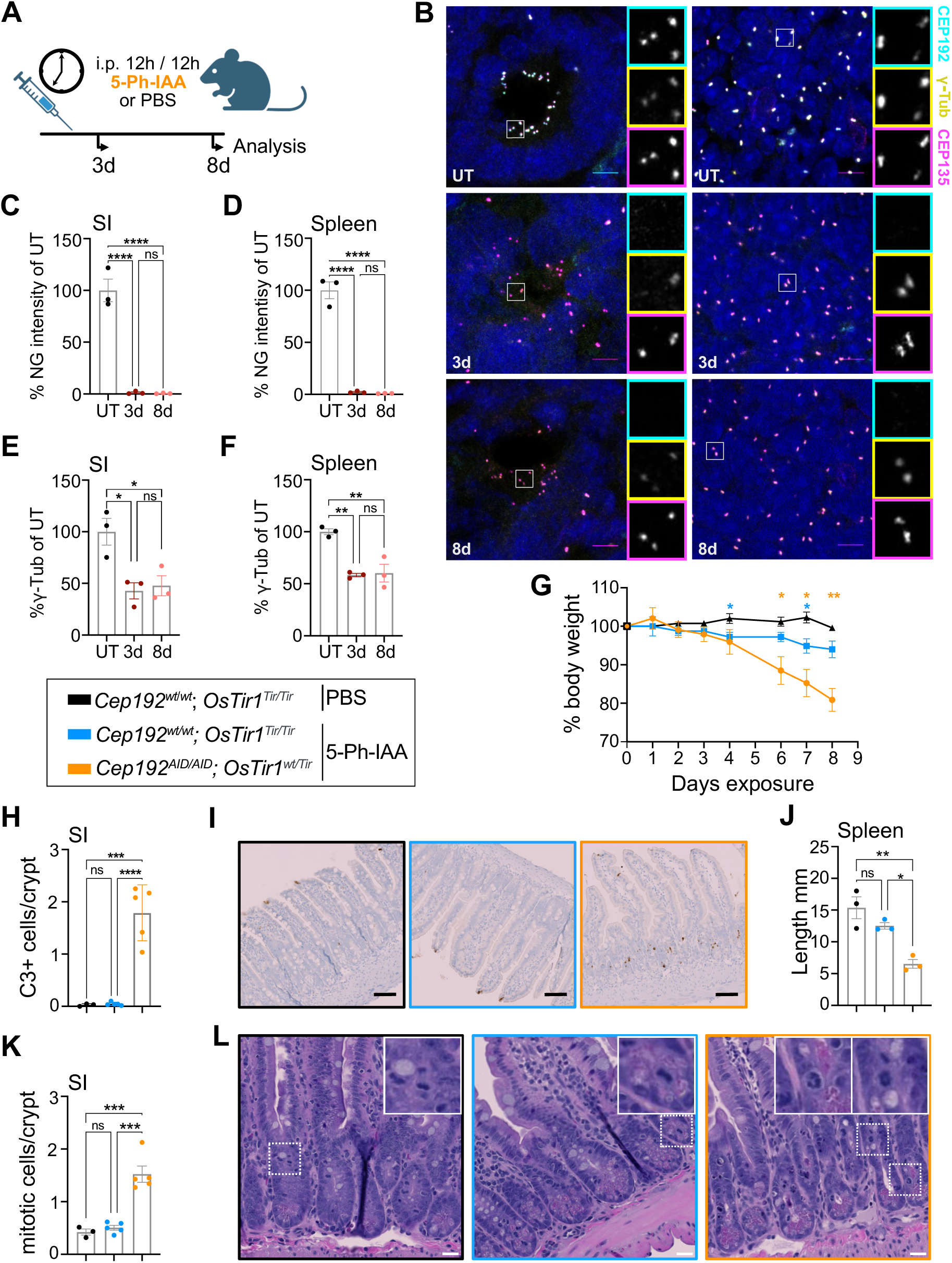
**Chronic degradation of CEP192^AID^ causes cell death in the gastrointestinal tract and weight loss.** (A) Schematic illustration of the experiment designed to test the effects of prolonged CEP192 depletion in mice. Mice were i.p. injected with PBS or 5-Ph-IAA every 12h and analyzed after 3 and 8 days. (B) Representative confocal immunofluorescence images of the small intestine (SI) and the spleen of *Cep192^AID/AID^*; *OsTir1*^wt/*Tir*^ mice left untreated (UT) or subjected to repeated 5-Ph-IAA injections for 3 or 8 days. CEP192^AID^ signal (cyan), γ-tubulin (yellow), and CEP135 (magenta); scale bars 5 µm. (C-D) Graphs showing the CEP192^AID^ signal (NeonGreen, NG) in immunofluorescence images of the (C) SI and the (D) spleen relative to the untreated control (UT) at the indicated time points. (E-F) Quantifications of the γ-tubulin signal intensity in immunofluorescence images in the (E) SI and (F) the spleen after 3 and 8 days. (G) Bodyweight changes of *Cep192^AID/AID^*; *OsTir1*^wt/*Tir*^ mice repeatedly injected with 5-Ph-IAA over 8 days. N = 5 mice per genotype (8 days); N = 3 mice per genotype (3 days). (H-I) Cell death was measured by immunohistochemistry for cleaved caspase-3 (C3) in the crypts of the small intestine (SI). Mouse tissues were analysed 8 days after treatment. (H) Quantification of cleaved caspase-3 positive cells per crypt. (I) Representative images of the SI. Black box: *Cep192^wt/wt^*; *OsTir1^Tir/Tir^* mice injected with PBS; Blue box: *Cep192^wt/wt^*; *OsTir1^Tir/Tir^* mice injected with 5-Ph-IA: Orange box: *Cep192^AID^*^/*AID*^; *OsTir1^wt/^*^Tir^ mice injected with 5-Ph-IAA (orange box). Scale bars 100 µm. (J) Quantification of the spleen size after 8 days of repeated 5-Ph-IAA injections and in untreated *Cep192^AID/AID^*; *OsTir1*^wt/*Tir*^ control mice (black). N = 3 mice per group. (K) Graph showing the number of mitotic cells per crypt quantified in H&E-stained paraffin sections of the small intestine (SI) of untreated *Cep192^AID/AID^*; *OsTir1*^wt/*Tir*^ mice (black box) or from mice injected with 5-Ph-IAA for 8 days (*Cep192^wt/wt^*; *OsTir1^Tir^*^/*Tir*^ mice, blue box; *Cep192^AID/AID^*; *OsTir1*^wt/*Tir*^ orange box). (L) Representative H&E-stained paraffin sections as in (K). Insets highlight mitotic cells. Scale bars 50 µm. Data is shown as mean ± SEM. Statistical significance was measured using one-way ANOVA (C-E, H, J, K) with Sidak’s multiple comparisons test, or mixed-effects analysis with Dunnett’s multiple comparisons test. ns p≥0.05, * p<0.05, ** p<0.01, *** p<0.001, **** p<0.0001

## Discussion

Here, we show that the second-generation AID system is well suited for acute and long-term targeted protein degradation in live mice. The AID2 system has been used successfully in *C. elegans* and *D. melanogaster*, and a proof-of-principle study by Yesbolatova et al. showed that the AID2 system effectively degraded a fluorescent reporter in mice (*18*, *39*, *40*). Our data add to this by showing that the AID2 system can drive near-complete degradation of an essential, endogenously tagged protein in live mice. The first-generation AID system uses IAA, which is toxic *in vivo* (Fig. 1), limiting its applications to *ex vivo* studies with primary cells or cell lines (*10*, *19*). In contrast, 5-Ph-IAA is well tolerated at concentrations required for protein degradation, and we did not observe leaky degradation in the absence of 5-Ph-IAA, as described with the first-generation AID system (Fig. 1) (*37*).

Other systems for *in vivo* protein degradation in mice, namely dTag and PROTACs, depend on large, complex molecules. Previous studies showed that effective delivery of these compounds requires vehicle formulations that can cause toxicity or inflammation, particularly after repeated injections (*7*, *11*). An advantage of AID2 over these systems is that 5-Ph-IAA is a small molecule molecular glue that can be administered in PBS or saline. In contrast to the dTag or the first-generation AID system, we did not observe any toxicity or inflammation at the injection site for 5-Ph-IAA (*11*, *19*). Moreover, we show that repeated dosing over two weeks with a concentration of 5-Ph-IAA that induces efficient degradation does not affect the well-being of the animals (Fig. 4). Besides toxicity, another advantage of the AID2 system is the remarkably fast degradation dynamics (Fig. 3). Our *in vivo* time course shows that, depending on the tissue, 80-90% degradation can be achieved within 30 minutes. This makes this system suitable for studying fast processes such as mitosis or cell death. One limitation of the AID2 system compared to the dTag system or PROTACs is the requirement for more complex mouse genetics. While both AID2 and dTag rely on endogenous tagging of the gene of interest, the AID2 system also requires the expression of *OsTir1*. The conditional *OsTir1* allele, however, allows for tissue specific degradation, which is difficult to achieve with other systems.

The Depmap classifies CEP192 as a pan-essential protein and therefore, it is not possible to study its function in mice *in vivo* with knockout mouse models. Here, we explored CEP192 biology in different cell types using the AID2 system in primary cultures and live mice. In line with prior work, we found that CEP192 is essential for bipolar spindle assembly in MEFs (Fig. 2) (*30*, *32*). Following CEP192 degradation, MEFs formed monopolar spindles, aborted mitosis and became polyploid. Importantly, proliferating intestinal cells also need CEP192 for successful mitosis (Fig. 5). Degradation of CEP192 for 8 days caused gastrointestinal defects because of massive cell death in the crypts of the small and large intestines. Together with the increased mitotic index, this indicates that in the absence of CEP192, proliferating intestinal cells cannot complete mitosis and undergo cell death. Moreover, continuous depletion of CEP192 reduced the spleen size, suggesting that cells in this proliferative organ also die in the absence of CEP192, presumably because of errors in mitosis (Fig. 5). In line with previous work, short-term ablation of CEP192 in interphase MEFs had no impact on γ-tubulin, but it was decreased in mitotic cells (Fig. 2) (*27*, *30*). Sustained CEP192 degradation in the SI and spleen reduced γ-tubulin abundance at the centrosome, suggesting γ-tubulin recruitment is impaired in the absence of CEP192 *in vivo* (Fig. 6). We also addressed the unstudied role of CEP192 in the formation of primary and motile cilia (Fig. 3). Although CEP192 is found at centrosomes and newly born centrioles in MEFs and MCCs, we show that it is dispensable for ciliogenesis and cilia maintenance. In summary, our data indicate that the essential *in vivo* role of CEP192 is in mitotic spindle assembly, explaining why proliferating tissues exhibit a higher dependence on CEP192. The *OsTir1* allele we created is conditional, allowing expression to be driven by tissue-specific *Cre* activity. Conditional expression of *OsTir1* in less proliferative tissues is likely to overcome the limited tolerance to whole-body depletion of CEP192 and enable future investigation of CEP192 function in other cell types.

## Acknowledgements

We thank all current and former members of the Holland lab for helpful discussions. This work was funded by National Institutes of Health grants R01GM114119, R01GM133897 and R01CA266199 to AJH, an EMBO Postdoctoral fellowship (ALTF 1194-2020) to VCS, and a Damon Runyon Cancer Research Foundation Merk Fellowship (DRG-2478-22) to CEJ. Parts of the schematic illustrations were created with Biorender.com (agreement Nr. EG26R9MTEM).

## Author contributions

Conceptualization: AJH, VCS; Methodology: VCS, CDG, PMS; Investigation: MAS, VCS, DTG, CEJ, CDG; Formal analysis: VCS, DTG, CEJ; Writing – Original draft: VCS, CEJ; Writing – Review and Editing: AJH, VCS; Visualization: VCS, CEJ; Supervision: AJH; Project administration: AJH; Funding acquisition: AJH, VCS.

## Materials & Methods

### Mice

*Rosa26^LSL-OSTir1-F74G^* and *Cep192*^mAID-mNeonGreen^ mice were generated using CRISPR/Cas9 technology. Briefly, Cas9 protein (30ng/µl, PNABio), tracrRNA (0.6μM, Dharmacon), crRNA (0.6μM, IDT) and ssDNA oligonucleotide (10ng/µl, IDT) were mixed and diluted in RNase-free injection buffer (10 mM Tris-HCl, pH 7.4, 0.25 mM EDTA) and injected into the pronucleus of one-cell embryos and subsequently transplanted into pseudopregnant ICR females. The sgRNAs were chosen based on the location of the available PAM sites and a minimal number of predicted off-target sites according to crispor.tefor.net. Injections and transplantation were performed by the JHU Transgenic Core.

*Rosa26^LSL-OsTir1-F74G^* mice were generated by targeting the *Rosa26* locus as described before (*41*). The sgRNA and the targeting vector containing a lox-stop-lox cassette, codon-optimized *OsTir1* (*42*) harboring the F74G modification and a myc tag (Fig. 1C) were co-injected into B6SJL/F2 embryos.

sgRNA: 5’- actccagtctttctagaaga

Primers used for genotyping and sequencing to characterize the offspring:

ROSA-P1: 5’- AAA GTC GCT CTG AGT TGT TAT

ROSA-P2: 5’- GCG AAG AGT TTG TCC TCA ACC

ROSA-P3: 5’- GGA GCG GGA GAA ATG GAT ATG

Following one round of backcross to a C57B/6J wild type, *Rosa26^LSL-OsTir1-F74G^* were crossed to Sox2-Cre females ((*43*) (The Jackson Laboratory strain #008454) to achieve full body expression of OSTIR1 by excision of the lox-Stop-lox cassette (*Rosa26^OsTir1-F74G^)*.

*Cep192*^mAID-mNeonGreen^ mice were generated by co-injection of an sgRNA and a DNA donor construct to target the last exon (48) of *Cep192* (Fig. 1E). Offspring with correct insertion was crossed to C57BL/6J wild type for two generations and then intercrossed to generate homozygous animals.

sgRNA1: 5’- cgactaatcggtgaatccct DNA donor template: 5’-

AGATGAAGGCAAGAGTATTGCTATTCGACTAATCGGCGAGTCTCTTGGAAAAAGTGGAGGTGCTGGCTCTGGACCTGGCTCTGGAGCAGGCGGTGGCAGCGGGGATCCAGCAGACTCTAAAGAGAAGTCAGCTTGTCCTAAGGACCCTGCCAAGCCACCTGCGAAGGCCCAAGTAGTCGGTTGGCCACCCGTGCGCAGCTATAGGAAGAACGTGATGGTGTCCTGTCAGAAGTCTAGCGGGGGGCCCGAGGCTAGCGGCGGATCTTCCTCAAAGGGAGAAGAAGACAACATGGCCTCACTGCCCGCAACACACGAGCTGCATATTTTCGGAAGCATCAATGGGGTGGATTTCGACATGGTGGGCCAGGGGACTGGAAACCCAAATGATGGGTACGAGGAACTGAATCTGAAGTCAACCAAAGGCGACCTCCAGTTCAGCCCTTGGATTCTGGTGCCCCACATTGGCTATGGGTTTCATCAGTATCTGCCCTACCCTGACGGAATGTCCCCATTCCAGGCAGCTATGGTGGATGGATCTGGCTACCAGGTCCACAGGACCATGCAGTTTGAGGACGGGGCCAGTCTGACTGTGAACTACCGCTATACCTACGAGGGATCACATATCAAGGGCGAAGCACAGGTGAAAGGGACAGGATTCCCAGCTGATGGCCCC GTCATGACAAACTCTCTGACCGCCGCCGACTGGAGCCGGTCCAAGAAAACTTACCCTAACGATAAGACCATCATCTCTACCTTCAAGTGGAGTTATACCACAGGCAACGGGAAGCGGTACAGAAGCACAGCCCGAACTACCTATACTTTTGCTAAGCCCATGGCTGCAAACTATCTGAAAAATCAGCCTATGTACGTGTTCAGAAAGACCGAGCTGAAGCACTCCAAAACAGAACTGAATTTCAAGGAATGGCAGAAGGCTTTTACCGATGTGATGGGGATGGACGAACTGTATAAGTAACTAGAATACATATTGTGTAAATGACCTGCTTATAT

Primers used for genotyping and allele characterization:

F1: 5’- GGT GCA CTT CAG ACC AGA GG

R2: 5’- TGT AGA AAA ATG AGA TTG CCA AA

AID-R: 5’- TGA CAG GAC ACC ATC ACG TT

Mice were housed under standard conditions in an AAALAC-accredited facility. All animal experiments were approved by the Johns Hopkins University Institute Animal Care and Use Committee (MO21M300). Age-matched 2-5 months old mice were used in a balanced sex ratio in all experiments.

### 5-Ph-IAA treatment *in vivo*

For intraperitoneal injections, 5-Phenyl-1H-indole-3-acetic acid (5-Ph-IAA, #30-003, BioAcademia, Osaka, Japan) was dissolved in sterile PBS. Specifically, 10mg of 5-Ph-IAA were dissolved in 1 mL PBS under dropwise addition of NaOH. Once dissolved, the pH was adjusted to 7.4 using minute amounts of HCl and reconstituted with PBS to achieve a concentration of 1 mg/mL. The solution was sterile filtered using 0.2 µM syringe filters. Mice were injected intraperitoneally with 1 mg/kg or 5 mg/kg either once or repeatedly every 24 hours (Fig. 5) or every 12 hours (Fig. 6). Control mice were injected with PBS for repeat dose experiments or left untreated for single dosing.

### Blood and serum analysis

Complete blood cell counts were performed by the Johns Hopkins University Phenotyping and Pathology Core using a DEXX ProCyte Dx® Hematology Analyzer. The standard tox panel of serum parameters was analyzed by IDEXX BioAnalytics, North Grafton, MA, USA.

### Histopathological assessment

4 µm H&E-stained sections of paraffin-embedded liver tissue were used for histopathological assessment by a board certified veterinary pathologist at the Johns Hopkins University Phenotyping and Pathology core.

### Isolation and differentiation of mouse tracheal epithelial cells (mTECs)

mTECs were isolated, cultured and differentiated as previously described (You and Brody, 2013). In brief, tracheas were dissected and incubated in Pronase (Roche) at 4°C overnight. Following enzymatic and mechanical dissociation, the tracheal cells were seeded onto 0.4 µm Falcon transwell membranes (Corning). After 5 days of proliferation, the medium was removed from the apical chamber and the basal medium was changed to NuSerum medium, starting the air-liquid interphase culture (ALI day 0). Cells were allowed to differentiate up to 9 days. For experiments in Fig 3 (E, G-I), 1µM 5-Ph-IAA was added on ALI day 0. To assess cilia maintenance, the mTEC cultures were treated with 1µM 5-Ph-IAA on days 7-9. For the time course experiments, 1µM 5-Ph-IAA was added on ALI day 5 (Fig. 3F, S3D). The basal medium with or without 5-Ph-IAA was refreshed every 1-2 days.

### Generation and culture of mouse embryonic fibroblasts

E12.5-14.5 embryos were isolated as previously described (*44*). In brief, the embryo bodies were digested at 4°C overnight in 0.05% Trypsin-EDTA (Gibco, 25300062). After 5 minutes incubation at 37°C, the MEF cells were dissociated by pipetting. MEFs were cultured in DMEM (Corning, #10-017-CV) containing 10% FBS (Corning, 35-010-CV), penicillin, streptomycin (Gibco, #10378016), and 0.1mM β-mercaptoethanol at 37°C in 5% CO2 and 3% O2 atmosphere. To induce degradation, the MEFs were treated with 1µM 5-Ph-IAA in DMSO or DMSO (control) for the indicated time periods up to 5 hours. For MEF ciliation, media was removed, cells were washed with PBS, and then serum starvation media (normal MEF media except only 0.5% FBS) was added either with or without 1µM 5-Ph-IAA.

### Imaging

#### Immunofluorescence

Fresh tissue was embedded in TissueTek O.C.T. compound (Sakura Finetek), sectioned on a Leica CM1950 cryostat (20 µm) and collected on Superfrost Plus microscope slides (Thermo Fisher Scientific). To examine ciliogenesis, MEFs and mTECs on coverslips were fixed in 4% paraformaldehyde in PBS. Sectioned tissues and MEFs on coverslips for all other experiments were fixed first in 1.5% paraformaldehyde (Electron Microscopy Sciences, #15714) and then in -20°C cold methanol for 4 minutes each. Following blocking in 2.5% FBS, 200 mM glycine, and 0.1% Triton X-100 in PBS for 1 hour, the samples were incubated with the respective primary antibodies in the same buffer for 1 hour. The samples were washed three times with PBS + 0.5% TritonX-100 and incubated with the secondary antibodies (Invitrogen) and DAPI. Samples were washed three times and mounted in Prolong Gold Antifade (Life Technologies, #P36930). Primary antibodies used were CEP135 (rabbit polyclonal, homemade, Alexa555- conjugated, 1:500), γ-Tubulin (goat polyclonal, homemade, Alexa647-conjugated, 1:500), CEP164 (rabbit polyclonal, EMD Millipore Corp., ABE2621, 1:1000), mouse CEP192 (rabbit polyclonal, gift from Karen Oegema, 1:1000), FOXJ1 (mouse monoclonal, ThermoFisher #14- 9965-82, 1:1000), Acetyl-α-Tubulin (Rabbit polyclonal, Lys40 (D20G3), Cell Signaling #5335T, 1:1000), mouse anti-PCNT (1:250, BD Transduction Laboratories #611814), ARL13B (mouse monoclonal, N295B66, Antibodies Incorporated #75-287, 1:1000). Tissue sections and MEFs (primary cilia assessment) were imaged on an SP8 confocal microscope (Leica Microsystems) using a Leica 40× 1.30 NA oil objective at 0.2-0.5 μm z-sections. For degradation analysis, MEFs were imaged using a Deltavision Elite system (GE Healthcare) with an Olympus 60× 1.42 NA oil objective at 0.2 µm z-sections. mTECs were imaged on a Zeiss Axio Observer 7 inverted microscope with Slidebook 2023 software (3i—Intelligent, Imaging Innovations, Inc.), CSU-W1 (Yokogawa) T1 Super-Resolution Spinning Disk, and Prime 95B CMOS camera (Teledyne Photometrics) with a 63x plan-apochromat oil immersion objective with 1.4 NA.

### Live imaging

MEFs were seeded in separate 4-well Cellview cell culture dishes (Greiner) and live cell imaged on an SP8 confocal microscope at 37°C, 5% CO2. 5-Ph-IAA (1 µM) was added to at t=0 min and images were taken of every 5 minutes for 75 minutes Leica 63×, 1.40 NA oil objective with 0.5 µm z-sections.

To measure mitotic duration and fate, MEFs were imaged (brightfield) using an IncuCyte S3 (Sartorius) every 20 minutes for 36h. To induce degradation, 5-Ph-IAA (1 µM) was added at t=0 min. The time in the stills shown in Supplemental Fig. 2C was adjusted so that one frame before rounding up of the cell was set to timepoint 0 min.

### Image analysis

All imaging analysis was performed blinded using ImageJ (v2.1.0/1.53c, US National Institutes of Health, http://imagej.net). For tissue sections, immunofluorescence images were lightning processed using LAS X Software (Leica, v3.5.6.21594) and maximum intensity projections (16 bit) were generated using ImageJ. Centrioles of MEFs were assessed on deconvolved 2D maximum intensity projections (16 bit). For live imaging of CEP192^AID^ degradation in MEFs, the CEP192^AID^ (NeonGreen) signal was quantified per frame in maximum intensity projected movies. For fixed and live imaging, signal intensities of CEP192^AID^ (NeonGreen), CEP135 and γ-Tubulin were determined by drawing a circular region of interest (ROI_S_) around the centrosome, and a larger circular ROI (ROI_L_) around the ROI_S_. The signal in ROI_S_ was calculated using the formula I_S_ – [(I_L_ – I_S_)/(A_L_ − A_S_) × A_S_], where A is area and I is integrated intensity.

For IncuCyte live imaging, mitotic cells were followed over time to determine the fate and the duration of mitosis. The beginning of mitosis was defined as rounding up of the cell. The end of mitosis was given by one of three possible fates: (1) successful division with a visible metaphase plate or anaphase and re-adhering of two daughter cells, (2) re-adhering without division (abortive mitosis), or (3) cell death.

### Protein extraction and Immunoblotting

Snap-frozen tissues were homogenized in RIPA lysis buffer (150 mM NaCl, 50 mM Tris, 1% NP- 40, 0.5% sodium deoxycholate 1% SDS, 1 tablet EDTA-free protease inhibitors) and protein content was determined using a Bradford protein assay (Bio-Rad, #5000001). 50-100 µg protein was run on an SDS-PAGE and blotted using a wet-transfer system (Bio-Rad) to a nitrocellulose membrane (0.45 µm, Santa Cruz Biotechnology, # sc-3724). Transfer quality and total protein were assessed by Ponceau-S staining before the membranes were probed for Myc-tag (Abcam, #ab9106, 1:1000). Fluorophore-conjugated secondary antibodies were used for detection using the Licor Odyssey CLx system (anti-rabbit IgG, Thermo Fisher, SA5-35571, 1:10000).

### Statistical analysis

Statistical analysis was performed using GraphPad Prism (v9.0.0, GraphPad software, LLC.). Two-tailed, unpaired Student’s t-test was used for comparison of two groups. One-way ANOVA or two-way ANOVA with Sidak’s multiple comparisons test was used for three or more groups. Significance levels and tests performed are stated in the figure legends. In some graphs, only statistically significant results are indicated. “N” represents the number of animals and “n” refers to the number of cells analyzed per mouse. Each mouse is considered a biological replicate.

---

**Supplemental Figure 1:**
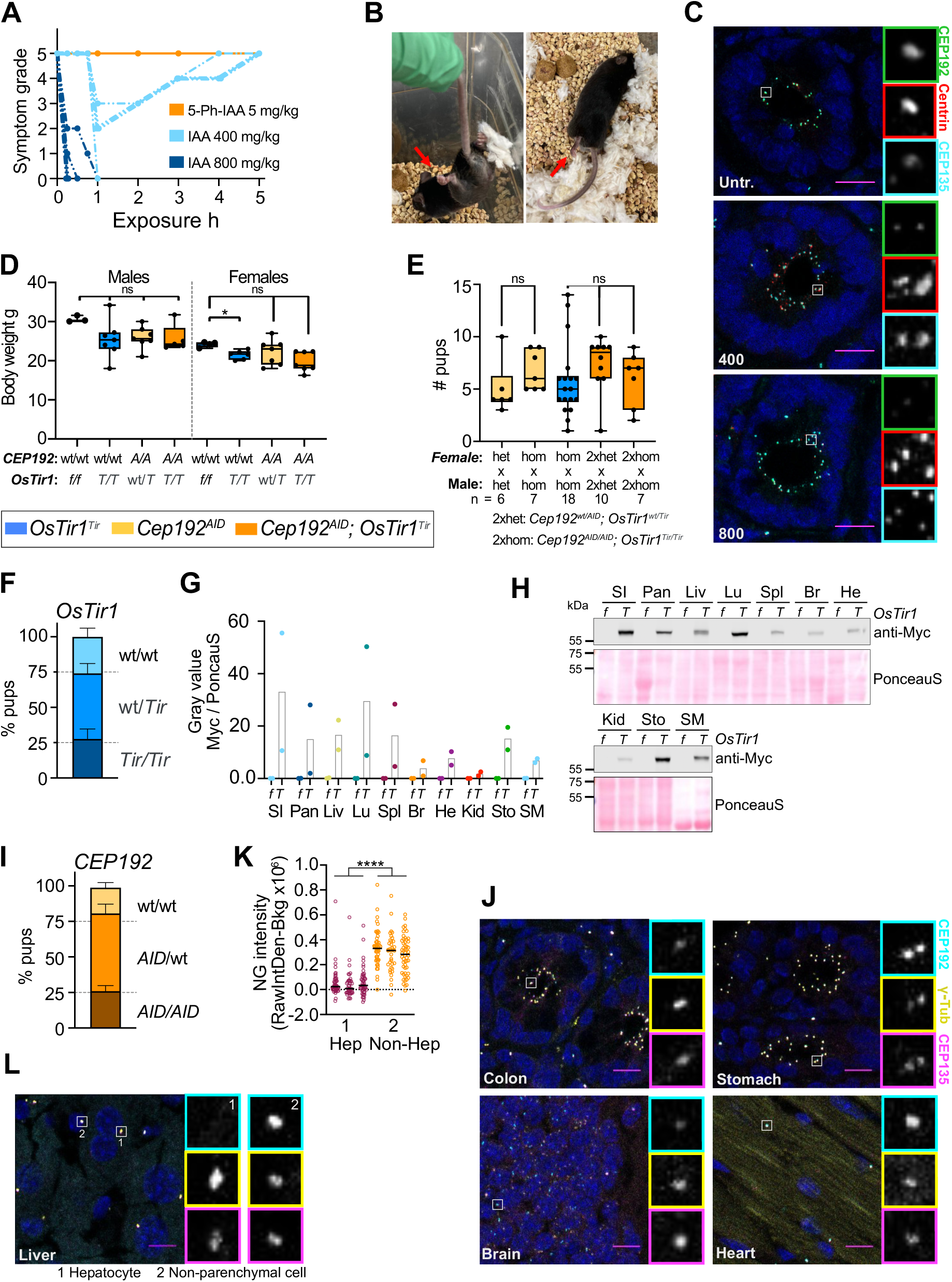
**Characterization of the novel *OsTir1^Tir^* and *Cep192^AID^* mice.** (A) Symptom grades of the individual *OsTir1^Tir/Tir^* mice injected either with 5mg/kg 5-Ph-IAA (N = 3), 400mg/kg IAA (N = 5), or 800mg/kg IAA (N = 5) as in Fig. 1B. Symptoms including spasms and paralysis were graded by severity over time; 0 = symptom-free, 5 = near complete paralysis, humane endpoint. (B) Pictures of *OsTir1^Tir/Tir^* mice 30min after injection of 800 mg/kg IAA. Red arrows mark cramping paws and limbs. (C) Representative confocal Immunofluorescence images showing CEP192^AID^ signal (NeonGreen; green), and immunostained Centrin (red) and CEP135 (cyan) in the SI of OSTIR1 expressing mice that were left untreated or injected with 400mg/kg for 5h or 800mg/kg IAA for 1h. Scale bar 10 µm. (D) Graph showing the body weight of 9-30 weeks old male and female mice of the indicated genotype combinations. *Cep192*: A/A – AID/AID, *OsTir1*: f/f – floxed/floxed, T/T – Tir/Tir. N = 3-8 mice per genotype. (E) Litter sizes of breedings with the indicated genotype combinations of homozygous (hom) or heterozygous (het) *Cep192^AID^* and *OsTir1^Tir^*alleles. n = 6-18 litters as noted in the figure. 2xhet: *Cep192^wt/AID^*; *OsTir1^wt/Tir^*. 2xhom: *Cep192^AID/AID^*; *OsTir1^Tir/Tir^*. (F) Genotype distribution of the offspring of *OsTir1^Tir^* heterozygous breedings. Dashed lines mark the expected Mendelian distribution. N = 9 litters. (G) Immunoblot probed with an antibody detecting OSTIR1-F74G-Myc in the indicated organs of an *OsTir1^f/f^ (f)* and an *OsTir1^Tir/Tir^ (T)* mouse. This experiment is a biological replicate of the immunoblot shown in Fig. 1D. SI – small intestine, Pan – Pancreas, Liv – Liver, Lu – Lung, Spl – Spleen, Br – Brain, He – Heart, Kid – Kidney, Sto – Stomach, SM – Skeletal Muscle. Ponceau- S-staining is shown as a reference for the amount of protein loaded. (H) Graph showing the genotype distribution of the offspring of heterozygous *Cep192^AID^* breedings. Dashed lines mark the expected Mendelian distribution. N = 11 litters. (I) Representative confocal immunofluorescence microscopy images of the indicated tissues as in (H) showing CEP192^AID^ signal (NeonGreeen, cyan), and immunostained γ-tubulin (yellow) and CEP135 (magenta). Scale bar 10 µm. (J) Graph showing the individual data points of the quantification in Fig. 1E. The CEP192^AID^ NeonGreen (NG) signal was measured as raw integrated density with background subtraction of N = 3 mice across tissues. SI – Small intestine, Col – Colon, Sto – Stomach, Pan – Pancreas, Liv – Liver, Lu – Lung, He – Heart, Spl – Spleen, Br – Brain. (K) Quantification and representative confocal immunofluorescence image of the CEP192^AID^ NeonGreen (NG) signal in hepatocytes (1) and non-parenchymal cells (2, non-Hep) in the liver as in (I-J). Scale bar 10 µm. Data is displayed as mean ± SEM. Statistical significance was determined by one-way ANOVA with Sidak’s multiple comparisons test. In (A), male and female groups were analyzed separately. ns p≥0.05, * p<0.05, **** p<0.0001.

**Supplemental Figure 2:**
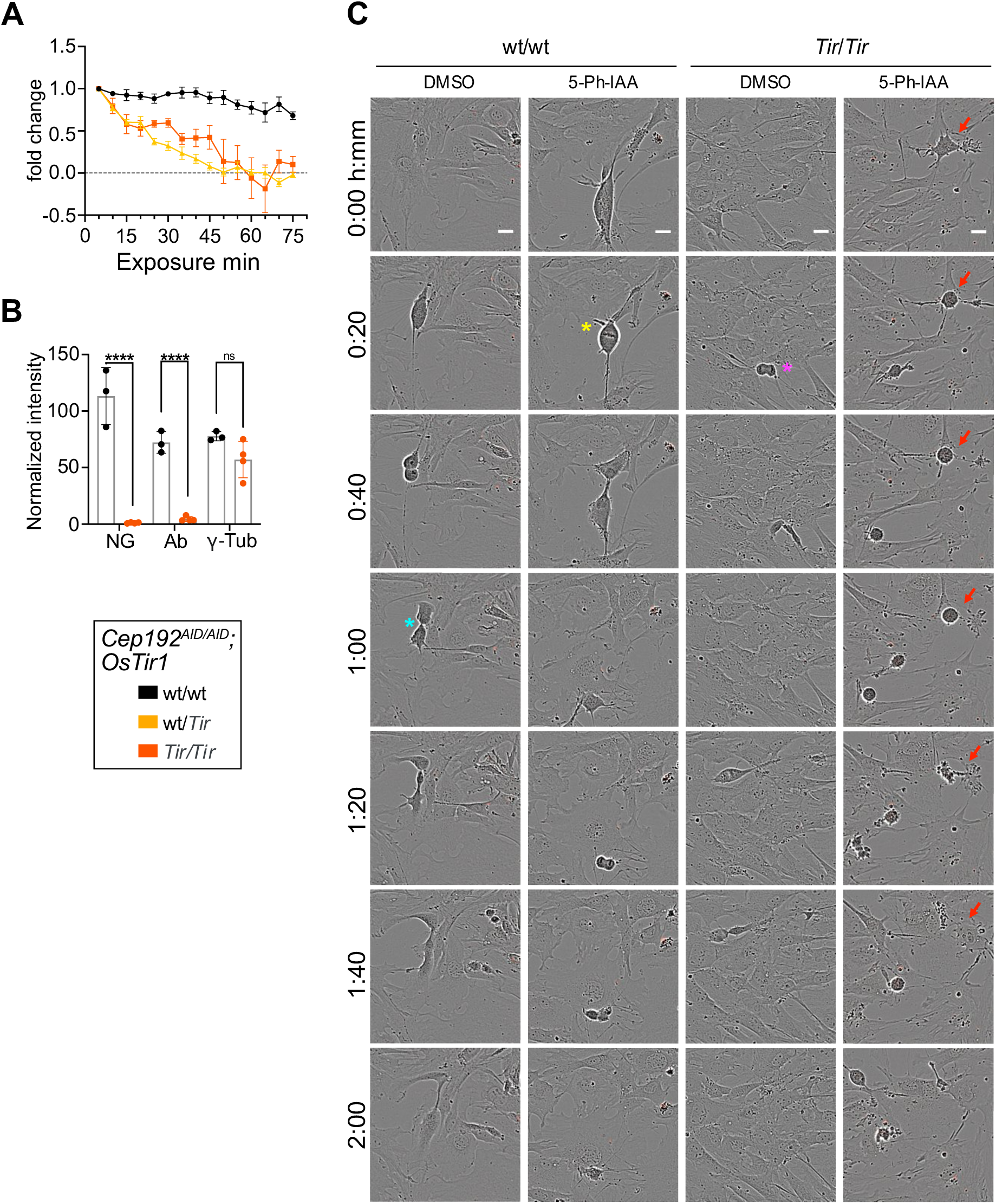
**Complete CEP192^AID^ in MEFs impairs mitosis.** (A) Graph showing the CEP192^AID^ NeonGreen signal intensity measured relative to timepoint t = 0 min in MEF lines of the indicated genotypes. 5-Ph-IAA was added at timepoint t = 0 min. *Cep192^wt^*^/*AID*^; *OsTir1^wt/wt^* N = 1 MEF lines, n = 4 cells; *Cep192^wt^*^/*AID*^; *OsTir1^wt/Tir^* N = 2, n = 3-8; *Cep192^wt^*^/*AID*^; *OsTir1^Tir/Tir^* N = 3, n = 3-11. (B) *Cep192^AID^*^/*AID*^; *OsTir1^Tir/Tir^* and *Cep192^AID^*^/*AID*^; *OsTir1^wt/wt^* MEF lines treated with 5-Ph-IAA for 5h were immunostained with an antibody raised against mouse CEP192 and an antibody recognizing γ-tubulin. Signal intensities of CEP192^AID^ (NeonGreen), and immunostained CEP192 and γ-tubulin were quantified relative to the DMSO condition. N = 3-4 MEF lines per genotype. (C) Representative live imaging brightfield stills of mitotic *Cep192^AID/AID^* MEFs with or without OSTIR1 treated with 5-Ph-IAA or DMSO. MEFs without OSTIR1 or 5-Ph-IAA undergo complete mitosis with visible metaphase plates (yellow *), anaphase (magenta *), and cytokinesis (cyan*). MEFs expressing OSTIR1 and treated with 5-Ph-IAA round-up and re-adhere to the plate without undergoing cell division (red arrow). All data is shown as mean ± SEM. Statistical significance was measured by two-way ANOVA ANOVA with Sidak’s multiple comparisons test (B). ns p≥0.05, **** p<0.0001.

**Supplemental Figure 3:**
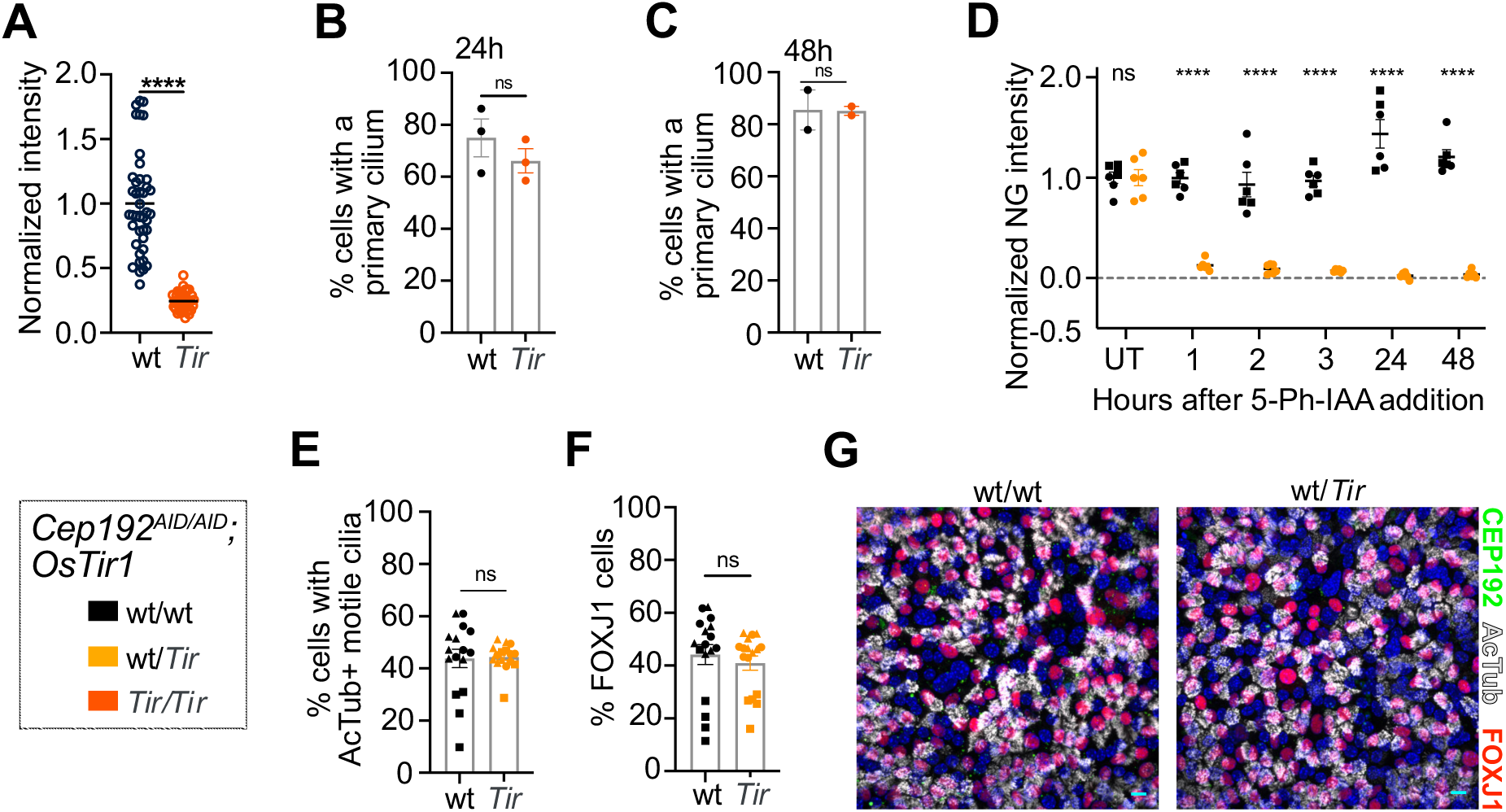
**CEP192 is not needed for primary or motile cilia maintenance** (A) Quantification of the NeonGreen signal of serum-starved *Cep192^AID^*^/*AID*^ MEFs with or without OSTIR1 normalized to the mean of *Cep192^AID^*^/*AID*^; *OsTir1^wt/wt^* MEFs. n = 39-42 cells per genotype. (B-C) Quantification of *Cep192^AID^*^/*AID*^ MEF cells wt/wt or wt/Tir for *OsTir1* with a primary cilium. Cells were serum starved for 24 h before 5-Ph-IAA treatment for (B) 24h or (C) 48h. N = 2 MEF lines per genotype from 1 or 2 separate passages. (D) Time course of 5-Ph-IAA treatment of mTEC cultures showing CEP192^AID^ (NeonGreen) intensity at the centrioles of non-multiciliated cells without or with OSTIR1 (wt/*Tir*) relative to untreated control. N = 2 mice per genotype indicated by symbol shape, n = 6 fields of view. (E-G) mTECs were cultured at an air-liquid-interphase (ALI) for 7 days to allow motile cilia formation. 5-Ph-IAA was added to the basal media for ALI days 7-9. (E) Quantification of the percentage of cells with motile cilia marked by acetylated tubulin (AcTub) and (F) the fraction of cells positive for the differentiation marker FOXJ1. N = 3 mice per genotype indicated by symbol shape, n = 6 fields of view. (G) Representative confocal images of mTEC cultures of the indicated genotypes expressing CEP192^AID^ (green) and immunostained for AcTub (gray) and FOXJ1 (red). Scale bars 10 µm. All data is shown as mean ± SEM. Statistical significance was measured by two -way ANOVA with Sidak’s multiple comparisons test (D), and two-tailed, unpaired Student’s t-test (A-C, E-F). ns p≥0.05, **** p<0.0001.

**Supplemental Figure 4:**
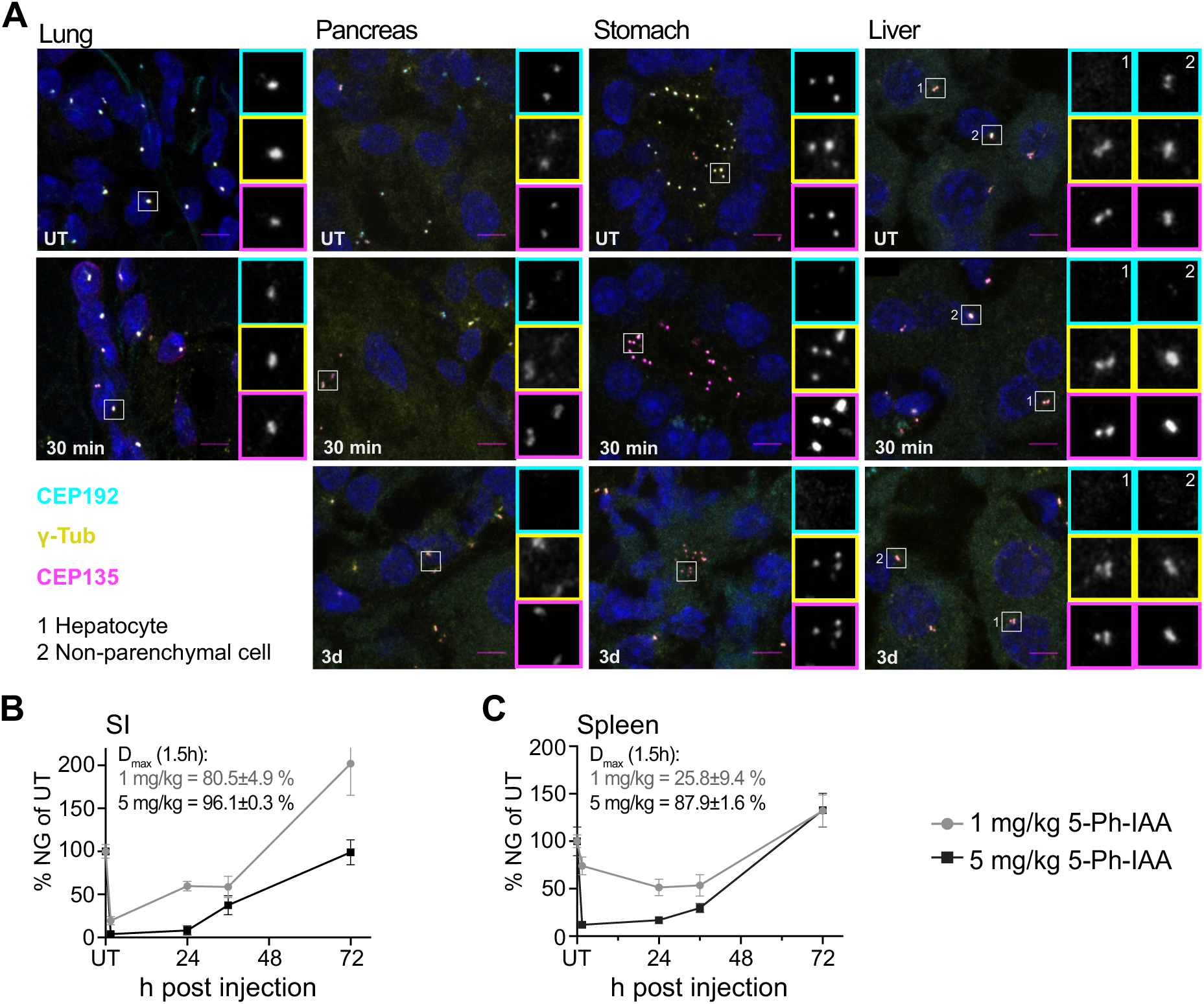
**Different doses of 5-Ph-IAA alter the degradation dynamics.** (A) Representative immunofluorescence images of organs from *Cep192^AID^*^/*AID*^ mice expressing OSTIR1 that were untreated (UT) or injected 5-Ph-IAA. Analysis was performed 30 min after 5-Ph-IAA administration or following repeat 5-Ph-IAA administration every 12 h for 3 days. CEP192^AID^ signal (cyan), and immunostained γ-tubulin (yellow) and CEP135 (magenta); scale bar 5 µm. (B-C) Quantification of the CEP192^AID^ NeonGreen (NG) signal relative to the untreated control (UT) of (B) the small intestine (SI) and (C) the spleen of mice injected with 1 mg/kg or 5 mg/kg 5-Ph-IAA at the indicated timepoints. All data is shown as mean ± SEM.

**Supplemental Figure 5:**
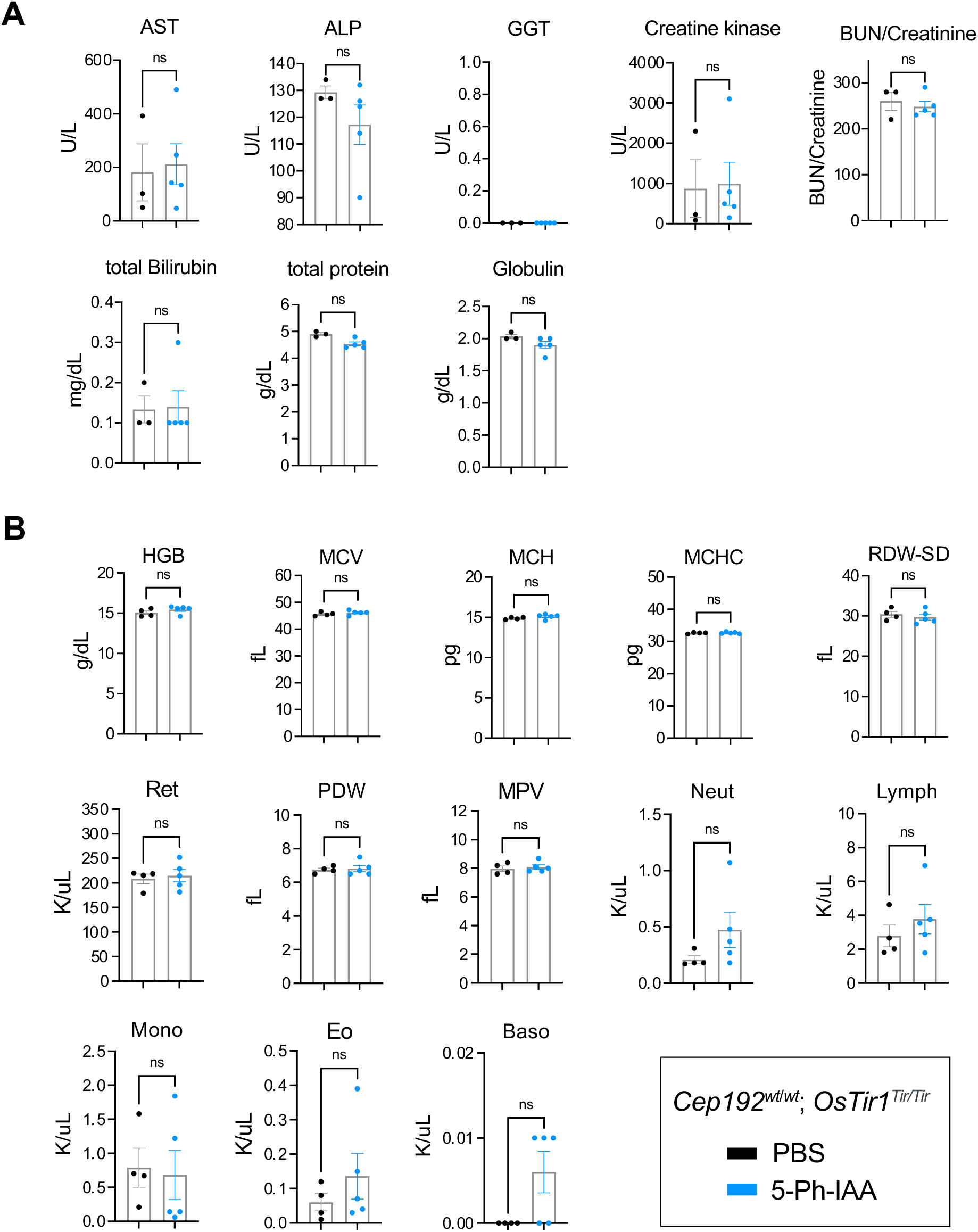
**Repeated dosing with 5-Ph-IAA has no impact on blood and serum parameters.** (A-B) Analysis of serum and blood parameters of *OsTir1^Tir/Tir^* mice treated with 5-Ph-IAA or PBS every 24h for 14 days. (A) Graphs showing the serum parameters. AST – Aspartate aminotransferase, ALP – Alkaline phosphatase, GGT – Gamma-glutamyltransferase, BUN – Blood urea nitrogen. (B) Graphs showing blood cell characterization. HGB – Hemoglobin, MCV – Mean corpuscular volume, MCH – Mean corpuscular hemoglobin, MCHC – Mean corpuscular hemoglobin concentration, RDW-SD – Red cell distribution width, Ret – Reticulocytes, PDW – Platelet distribution width, MPV – Mean platelet volume, Neut – Neutrophils, Lymph – Lymphocytes, Mono – Monocytes, Eo – Eosinophils, Baso – Basophils. Data is displayed as mean ± SEM. Statistical significance was assessed by two-tailed, unpaired Student’s t-test. ns p≥0.05.

**Supplemental Figure 6:**
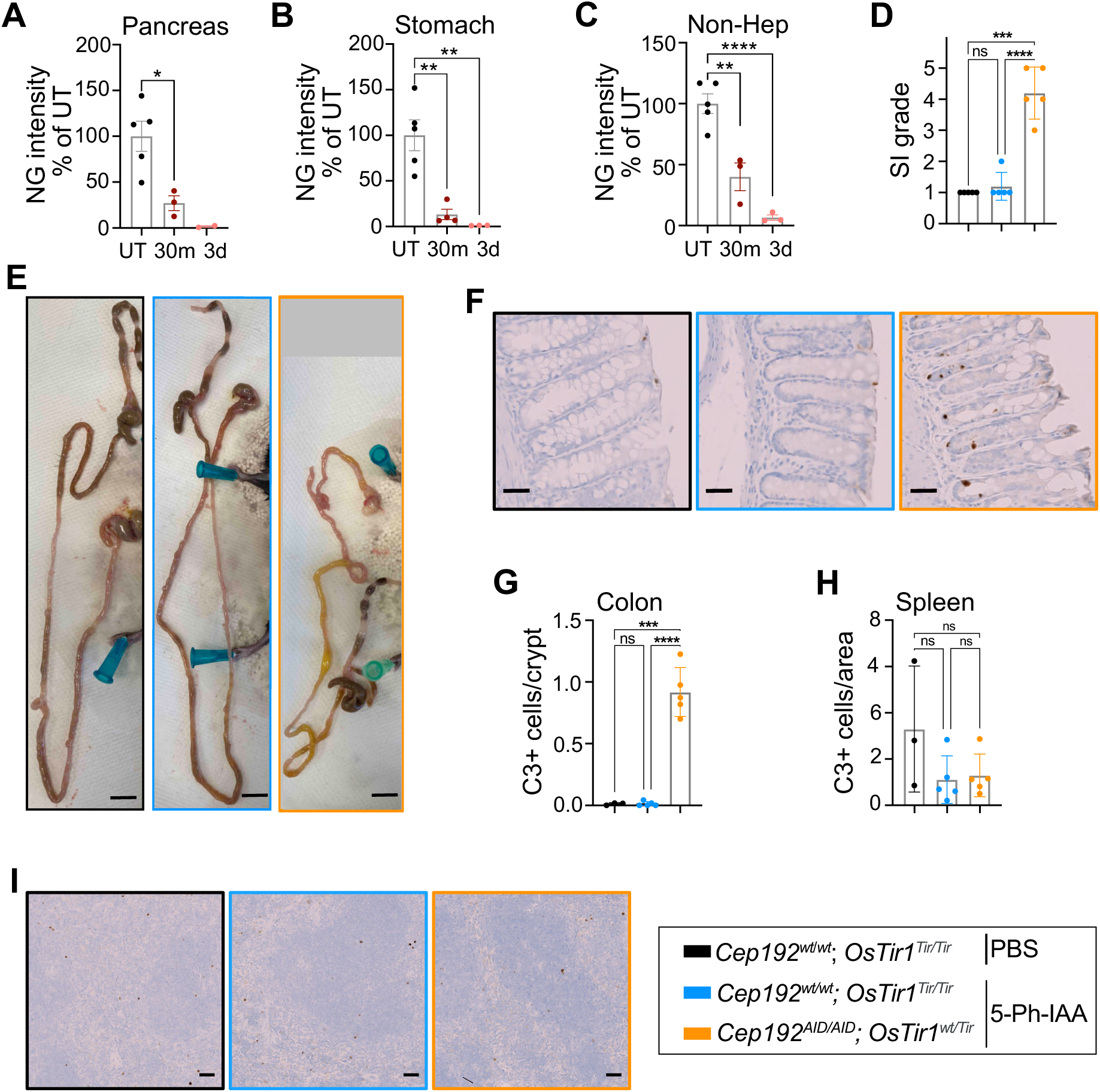
**Sustained degradation of CEP192^AID^ causes gastrointestinal symptoms and cell death.** (A-C) *Cep192^AID^*^/*AID*^ mice expressing OSTIR1 were injected with 5 mg/kg 5-Ph-IAA. Analysis performed 30 min after 5-Ph-IAA administration or following repeat 5-Ph-IAA administration every 12 h for 3 days. CEP192^AID^ signal intensity (NeonGreen, NG) was quantified in fluorescence microscopy images relative to the untreated (UT) control in (A) the pancreas (N = 2-5), (B) the stomach (N = 3-5) and (C) in non-parenchymal liver cells (N = 3-5) at the indicated timepoints. (D) Graph showing the severity of the gastrointestinal symptoms in mice after 8 days of repeated 5-Ph-IAA or PBS injections. Symptoms were graded 1-5 based on stool consistency and color in the small intestine. 5 = normal stool; 1 = watery liquid, yellow stool. (E) Example images for (D) showing SI grade 1 and 5 symptoms. Images from *Cep192^wt/wt^*; *OsTir1^Tir/Tir^* mice treated with PBS (black box) or 5-Ph-IAA (blue box) and *Cep192^AID^*^/*AID*^; *OsTir1^wt/Tir^* exposed to 5-Ph-IAA (orange box). Scale bar 1cm. (F) Cell death in the crypts of the colon was measured using immunohistochemistry for cleaved caspase-3 (C3). Representative images of the colon of *Cep192^wt/wt^*; *OsTir1^Tir/Tir^* mice treated with PBS (black box) or 5-Ph-IAA (blue box), and *Cep192^AID^*^/*AID*^; *OsTir1^wt/Tir^* mice injected with 5-Ph-IAA (orange box) for 8 days. Scale bars 100 µm. (G) (G) Graph showing the number of cleaved caspase-3 positive cells per crypt. (H) Quantification of cleaved caspase-3 positive cells in the spleen per area. Cell death was measured by immunohistochemistry for cleaved caspase-3 (C3). (I) Representative images of the spleen of *Cep192^wt/wt^*; *OsTir1^Tir/Tir^* mice injected with PBS (black box) or 5-Ph-IAA (blue box), and *Cep192^AID^*^/*AID*^; *OsTir1^wt/Tir^* exposed to 5-Ph-IAA (orange box) for 8 days. Scale bars 100 µm. All data is shown as mean ± SEM. Statistical significance was measured using one-way ANOVA with Sidak’s multiple comparisons test. Of note, no statistical analysis was performed for (A) 3d, since this timepoint includes only N = 2 mice. ns p≥0.05, * p<0.05, ** p<0.01, *** p<0.001, **** p<0.0001.

